# PSite: inference of read-specific P-site offsets for ribosomal footprints

**DOI:** 10.1101/2023.06.27.546788

**Authors:** Yue Chang, Tianyu Lei, Hong Zhang

## Abstract

**Summary:** Ribosome profiling is a powerful method for global survey of ribosomal footprints. Inferring the offsets of footprint 5’ ends to the ribosomal P-site is essential to pinpoint codons translated by ribosomes. By convention, global or read length-specific P-site offsets are estimated by inspecting the distribution of ribosome footprints around the annotated start or stop codons. However, actual offsets might be different even for footprints of the same length due to the influence of sequence context and the cutting bias of endoribonucleases. To address this issue, we present PSite, a python package for inferring read-specific P-site offsets using a gradient boosting trees model. PSite assigned more reads to the correct reading frame than conventional methods and improved the prediction of translated ORFs by existing software. Besides, PSite is robust to ribosome profiling datasets of varying quality or using endonucleases with cutting bias for digestion.

**Availability and implementation:** https://github.com/gxelab/psite.

## 1 Introduction

Ribosome profiling (Ribo-Seq) and derived methods enable genome-wide investigation of translation regulation from different facets (Ingolia, et al., 2019; Wang, et al., 2020). During Ribo-Seq, translating ribosomes are stalled by elongation inhibitors, and mRNA regions unoccupied by ribosomes are digested with Ribonuclease I (RNase I) or other endoribonucleases, leaving only ribosome-protected fragments (RPFs). By enriching and sequencing RPFs, a snapshot of translational activities in the cells can be obtained (Ingolia, et al., 2019). Although RPFs are usually 30 nt in length, each is derived from a ribosome translating a single codon. Therefore, an essential step in analyzing Ribo-Seq data is determining the offset between the 5’ ends of RPFs and codons being translated at the ribosomal P-site.

P-site offsets are usually estimated in a read length-specific manner by inspecting the distribution of the distance from RPF 5’ ends to annotated start or stop codons as in plastid (Dunn and Weissman, 2016) or RiboWaltz (Lauria, et al., 2018). However, these approaches ignore the influence of read-specific sequence context and the cutting bias of endonucleases used to digest unprotected RNA regions, which might hamper the accurate assignment of RPFs to P-site codons and downstream analyses. Although it is possible to estimate read-specific offsets with tree-based models (Fang, et al., 2018; VanInsberghe, et al., 2021), no standalone implementation is available for custom downstream analyses, and whether this approach can be generalized to different Ribo-Seq datasets is unclear. Therefore, we developed PSite, a python package for model-based inference of footprint-specific P-site offsets. We demonstrated that PSite is superior to conventional methods and robust to different Ribo-Seq datasets.

## 2 Implementation

PSite was implemented in Python 3 and required the alignment of RPFs against the reference transcriptome to train a gradient boosting trees (GBT) classifier. We used the GBT model because it is built by successively adding weak learners and is less prone to overfitting (Friedman, 2001). To exclude reads mapped to multiple genomic positions, we recommend mapping RPFs to the reference genome and then deriving the transcriptome alignment with STAR (Dobin, et al., 2013). We used RPFs whose 5’ ends are 10∼14 nt away from annotated start codons or 13∼17 nt away from annotated stop codons for training. As typical P-site offsets are about 12nt, RPF 5’ ends in the above range are most likely generated by ribosomes translating the start codon or the codon before the stop codon rather than flanking ones. For each RPF, read length and the six nucleotides flanking the 5’ end were used as features for prediction. Then the trained model can be used to predict offsets for RPFs mapped to the reference genome or transcriptome. PSite outputs bam files with a “PS” tag, which is convenient for custom downstream analyses. As a proof of concept, we implemented a submodule to calculate genome-wide P-site coverage employing the “PS” tag. Besides, PSite can also produce alignments of the P-site nucleotide only.

## 3 Results and discussion

To test whether PSite could effectively recover P-site offsets of RPFs, we examine the proportion of reads assigned to the correct reading frame for RPFs within annotated CDSs. In comparison, we also predicted the P-sites of RPFs using a fixed global offset of 12 nt (“Fixed”) or read length-specific offsets that were estimated from the peak distance of 5’ ends to annotated start and stop codons (“Peak”). We used the following datasets for testing: i) regular RNase I data where most RPF 5’ ends are in frame 0 (Patraquim, et al., 2022); ii) RNase I data where RPF 5’ ends are less than 50% in all three frames (Kronja, et al., 2014); iii) micrococcal nuclease (MNase) data (Zhang, et al., 2018). While all three methods performed well on regular RNase I data, only PSite assigned >50% RPFs to frame 0 for the other two datasets (Fig. 1A and B).

**Fig. 1.**
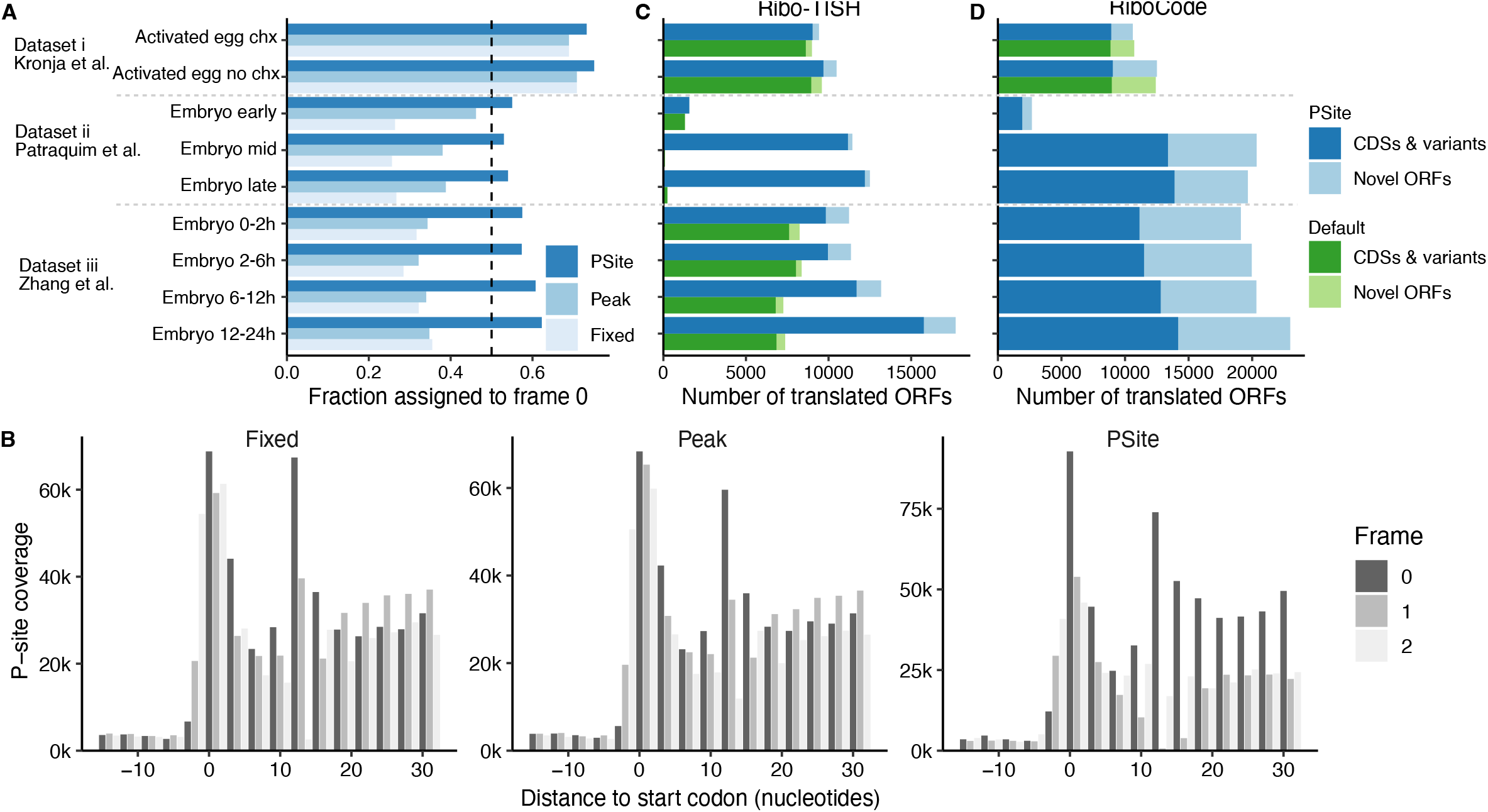
Performance of PSite. (**A**) Fraction of RPFs assigned to frame 0 by different offset estimation methods. (**B**) Metagene profiles of P-site coverage around CDS start codons for 0-2h embryos. (**C-D**) Number of translated ORFs predicted by either Ribo-TISH (**C**) or RiboCode (**D**) using the default or PSite pipeline.

To determine whether offsets inferred by PSite are biologically meaningful, we predicted translated ORFs using either the conventional or PSite-assisted pipeline. By default, both methods predict offsets internally with the “Peak” method. Compared to the default procedure, Ribo-TISH (Zhang, et al., 2017) predicted significantly more translated ORFs with the PSite pipeline (Wilcoxon signed-rank test, *P* = 0.004). In addition, most predicted ORFs are annotated CDSs and N-terminal extension or truncation variants (Fig. 1C). RiboCode (Xiao, et al., 2018) has a requirement on the minimum proportion of RPF 5’ ends in frame 0 and cannot run with the MNase or irregular RNase I datasets. However, with the PSite pipeline, thousands of translated ORFs are predicted, and the proportions of annotated CDSs or CDS variants are similar to those in the regular RNase I dataset (Fig. 1D).

In summary, PSite could assign most RPFs to the correct reading frame and is robust across different Ribo-Seq datasets. With other software, PSite allows flexible downstream analyses such as translated ORF prediction with improved performance.

## Data availability

The source of public data and code underlying analyses in this manuscript are available from the associated GitHub repository: https://github.com/gxelab/psite_analysis.

